# Structural characterization of functionally important chloride binding sites in the marine *Vibrio* alkaline phosphatase

**DOI:** 10.1101/2022.07.20.500776

**Authors:** Sigurbjörn Markússon, Jens G. Hjörleifsson, Petri Kursula, Bjarni Ásgeirsson

## Abstract

Enzyme stability and function can be affected by various environmental factors, such as temperature, pH and ionic strength. Enzymes that are located outside the relatively unchanging environment of the cytosol, such as those residing in the periplasmic space of bacteria or extracellularly secreted, are challenged by more fluctuations in the aqueous medium. Bacterial alkaline phosphatases (APs) are generally affected by ionic strength of the medium, but this varies substantially between species. An AP from the marine bacterium *Vibrio splendidus* (VAP) shows complex pH-dependent activation and stabilization in the 0 – 1.0 M range of halogen salts and has been hypothesized to specifically bind chloride anions. Here, using X-ray crystallography and anomalous scattering, we have located two chloride binding sites in the structure of VAP, one in the active site and another one at a peripheral site. Further characterization of the binding sites using site-directed mutagenesis and small angle X-ray scattering (SAXS) showed that upon binding of chloride to the peripheral site, structural dynamics decreased locally, resulting in thermal stabilization of the VAP active conformation. Binding of the chloride ion in the active site did not displace the bound inorganic phosphate product, but it may promote product release by facilitating rotational stabilization of the substrate-binding Arg129. Overall, these results reveal the complex nature and dynamics of chloride binding to enzymes through long-range modulation of electronic potential in the vicinity of the active site, resulting in increased catalytic efficiency and stability.

## Introduction

Alkaline phosphatase (AP) is a well-characterized model enzyme present in many species from bacteria to humans ^*1*^. While the activity of mammalian APs is mostly unaffected by ionic strength ^*2–4*^, salt concentration is known to affect the catalysis of bacterial APs, in particular anions ^*5–7*^. Since all the important active site features and the mechanism for hydrolysis are believed to be the same in all APs, it is not clear what causes the differences in salt activation between APs. Differences in environmental factors may have played a role through evolutionary time. The periplasmic space of Gram-negative bacteria, where AP is located, constitutes up to 40% of the total cell volume and is the location of several other enzymes. The outer membrane is relatively permeable to small molecules, and therefore, the periplasmic interior is highly exposed to fluctuations in the external environment ^*8*^. In cold-water marine bacteria, enzymes present in the periplasmic space face the challenge of adapting to a highly saline environment and low temperature. The salt and pH conditions are broadly the same in the periplasm as in the ocean, ~ 0.55 M chloride, ~ 0.47 M sodium with pH 7.6-8.4 ^*9–12*^.

Alkaline phosphatase from the marine bacterium *Vibrio splendidus* (VAP) is a cold-active enzyme located in the periplasmic space and one of the most catalytically active APs known ^*13*^. VAP activity is greatly affected by sodium chloride, reaching a maximum at salt concentrations close to oceanic salinity ^*14*^. Structurally, it is one of the largest APs, including several genetic inserts compared with related enzymes. One of the inserts forms a large interface loop that extends from each monomer to embrace the other subunit in the dimer close to the active site. This close association has an undefined role in promoting the high catalytic efficiency of this homodimeric enzyme ^*15*^. Two additional inserts are part of the so-called crown-domain, while the fourth insert forms a peripheral three-turn helix with an unknown function ^*16*^.

The salt activation of VAP is governed by the anion and accompanied by a large increase in thermal stability (~30 °C increase in T50%with 0.5 M NaCl) ^*13, 14*^, whereas in the absence of NaCl, the enzyme inactivates rapidly at room temperature. The stimulatory effect of anions on activity and stability was maximal at pH of 7-8, but absent at pH >10, and reached a plateau at ~ 0.5 M NaCl ^*14*^. These results fit well with pH dependent activation of *V. alginolyticus* AP reported by Hayashi et al. (1973) ^*17*^, an AP variant that is 90% sequence homologous to VAP. Some other enzymes from marine bacteria have been found to be stabilized and/or activated by chloride ions *in vitro*, and in more extreme cases, found to depend on chloride for function. *Vv*PlpA phospholipase from *Vibrio vulnificus* serves as an example of the latter, where an incomplete Ser-His-Gly catalytic triad was rescued by the binding of chloride, which in turn took the place of the traditional aspartic acid of the catalytic triad ^*18*^. The comparison of endonuclease I from the cold-adapted *Vibrio salmonicida* or mesophilic *Vibrio cholera* strains showed large differences in NaCl requirements, as well as dissimilar temperature stability, optimum pH and catalytic efficiency ^*19, 20*^. The optimal NaCl concentrations for the two variants corresponded well with the salinities in the natural habitats of the two host bacteria. The *E. coli* AP enzyme is also stimulated by salt ions despite not coming from a marine environment, yet the effect is not pH-dependent ^*7*^. Using fast-flow kinetics and inhibitor titrations it was shown that guanidinium chloride enhanced activity of the *E. coli* AP by accelerating the rate-limiting dissociation of the noncovalent E•P complex and by abolishing negative co-operativity that depended on the cooperation of the two subunits ^*21*^.

How chloride ions promote the catalytic process is not at all clear. For APs, the rate-limiting step at neutral to alkaline pH is the release of inorganic phosphate ^*6, 22,23*^, and the ionic effect might simply involve charge complementation of chloride ions with the binding site of the phosphate product, thus promoting its release. However, the lack of a similar effect with mammalian APs would seem to exclude such direct involvement in the reaction mechanism. Another possibility could involve consideration of dynamics, predicting a conformational change as being rate-limiting during the catalytic cycle. An early proposal of a half-of-sites mechanism (negative cooperativity), where the active sites alternate between high and low affinity for substrate/product ^*24–27*^, is still controversial.

In the current study, the structural basis of the anionic activation and stabilization of VAP was examined. VAP co-crystallization with chloride or bromide, together with anomalous mapping, allowed for localization of two chloride binding sites. Along with SAXS, mutagenesis and kinetical data, the crystal structures support a model, in which chloride binding in a solvent-exposed site distant from the active site facilitates stabilization by reducing thermal motions, and that diffusion-dependent active site binding facilitates catalytic rate increase through modulation of active site mobility and electrostatic potential.

## Materials and methods

### Materials

Chemicals were obtained from Sigma-Aldrich (Schnelldorf, Germany) or Merck (Darmstadt, Germany), unless stated otherwise. L-rhamnose and isopropyl β-D-1-thiogalactopyranoside (IPTG) were obtained from AppliChem (Germany). Bacto yeast extract and Bacto tryptone were purchased from Becton Dickinson and Company (France). Triton X-100 was obtained from BDH chemicals (England) and chloramphenicol from Ampresco (USA). Para-nitro-phenyl phosphate (pNPP) sodium salt was obtained from Thermo Scientific.

### Mutagenesis

Site-directed mutagenesis was performed using the QuikChange method as previously described ^*28*^. All constructs were confirmed by Sanger sequencing (Genewiz, Leibzig Germany).

### Protein expression and purification

Wild-type VAP (UNIP: Q93P54) and mutants, containing the original N-terminal periplasm targeting sequence and a C-terminal StrepTagll sequence ^*14*^, were expressed from a pET11a plasmid in Lemo(21)DE3 competent *E. coli* cells ^*29*^ and purified on StrepTactin column as previously described ^*14*^. The enzyme variants were subjected to further purification on a HiPrep Q XL 16/10 anion exchange column (GE Healthcare) pre-equilibrated in 20 mM Tris pH 8.0, 10 mM MgCl_2_, 15% (v/v) ethylene glycol at 4°C. Bound protein was eluted using a 0-100% linear gradient against the same buffer supplemented with 1 M NaCl. To avoid heterogeneity between different preparations in activity assays and crystallization, purified VAP was dialyzed against 20 mM Tris pH 8.0, 10 mM MgCl_2_, 300 mM NaCl and 15% (v/v) ethylene glycol using 3.5-kDa molecular weight cut-off (MWCO) dialysis membranes, at 4-8 °C overnight. Following dialysis, purified protein was concentrated to 10-25 mg/mL using 30 kDa MWCO Amicon^®^ Ultra 2-mL centrifugal filters (Merk, Germany), snap-frozen in liquid N_2_ and stored at −20 °C.

### Protein crystallization and structure refinement

We have previously crystallized the wild-type VAP construct (PDB:6T26), and the crystal structure contained tightly bound phosphate in both active sites, likely originating from overexpression in *E. coli* ^*30*^. In an attempt to remove the bound phosphate for co-crystallization with chloride, the enzyme was subjected to extensive dialysis (~48 h at 4 °C, with frequent renewal of buffer) against 20 mM CAPS pH 10, 500 mM NaCl, 10 mM MgCl_2_ and 15% (v/v) ethylene glycol, as VAP affinity towards phosphate is reduced by an order of magnitude at alkaline pH ^*14*^. The protein buffer was then exchanged back to the pH 8.0 buffer through dilution prior to concentration and crystallization.

Crystallization of wild-type VAP was carried out using hanging drop vapor diffusion at 20°C, in 24-well hanging drop plates (Molecular Dimensions). For co-crystallization with 0.5 M NaCl, 1.5 μL of 13.4 mg/mL wild-type VAP was mixed with 1.5 μL of the precipitant solution (26% PEG3350, 0.5 M NaCl, 1.0 M Tris pH 7.0) and placed over 1000 μL of the same solution. Rod shaped crystals with approximate dimensions of 200×100×50 μm^3^ grew over-night.

When conducting co-crystallization with 1.0 M NaCl, crystals grew in 3 μL equal volume drops in 28% PEG3350, 1.0 M NaCl, 0.1 M Tris pH 7.0 (1000 μL reservoir) from 13.4 mg/mL, protein after nucleation had been induced *via* streak seeding from crystals grown in 28% PEG3350, 0.8 M NaCl, 0.1 M Tris pH 7.0. Streak seeding was performed using a single pony hair, once the protein in the receiving crystallization drop had reached supersaturation, as indicated by phase separation. A single crystal with approximate dimensions of 1000×300×200 μm^3^ grew in two days after seeding.

VAP was co-crystallized with bromide at 13.4 mg/mL protein in 24% PEG3350, 1.0 M KBr and 0.1 M Tris pH 7.0, after inducing nucleation with seeds streaked from non-diffracting crystals grown in 24% PEG3350, 0.4 M KBr and 0.1 M Tris pH 7.0, using the procedure outlined above. Rod-shaped crystals with approximate dimensions of 200×50×50 μm^3^ grew in around 2 days after seed introduction.

VAP was co-crystallized with HEPES at 10 mg/mL protein in 23% PEG3350, 0.4 M NaCl and 0.1 M HEPES pH 7.5. As initial HEPES-bound crystals proved to be non-diffractive, subsequent crystals were dehydrated prior to freezing. Once crystals had grown, the cover slip containing the crystallization drop was moved onto a reservoir containing the same solution, but with the concentration of PEG3350 increased by 2% and incubated for ~1 h at 20 °C, before repeating the process in a stepwise manner until the final precipitant concentration reached 30%. All crystals were cryoprotected in their respective reservoir solutions supplemented with 25% (v/v) ethylene glycol and plunge-frozen in liquid N2 before shipping to synchrotron beamline facilities for diffraction data collection.

Diffraction data were collected on the P11 and P14 beamlines at the PETRA III synchrotron (DESY, Hamburg, Germany) ^*31*^, and on the BioMAX beamline ^*32*^ at the MaxIV synchrotron (Lund, Sweden). For VAP co-crystallized with NaCl, high-resolution data were collected at the standard beamline wavelengths (0.967-1.033 Å/12.0-12.7 keV), and additional, lower-resolution, anomalous datasets collected at 6 keV (2.066 Å) from the same crystals to obtain sufficient anomalous signal from chloride (K-edge of chlorine: 2.82 keV) for anomalous mapping. For the confirmation of halogen binding to VAP, diffraction data were collected from bromide co-crystals at 13.7 keV (0.905 Å), slightly above the theoretical K-edge of bromine (13.47 keV), to locate bound bromide *via* anomalous mapping. Diffraction data were processed in XDS and XSCALE ^*33*^. For the longer-wavelength chloride datasets, strict absorption correction (Friedel pairs treated as different reflections during estimation of absorption correction factors) was employed during processing to maximize the anomalous signal. Initial estimation of diffraction data quality was carried out in XTRIAGE ^*34*^, and phases solved using molecular replacement (MR), in phaser.MRage ^*35*^, with the previously published crystal structure of VAP (PDB: 3E2D) as the reference model. Model refinement was carried out in phenix.refine ^*36*^ within the Phenix software suite (v. dev-3958), and manual model building in COOT ^*37*^. Model validation was performed using MolProbity ^*38*^. Diffraction data collection and refinement statistics are shown in Table 1.

**Table 1:**
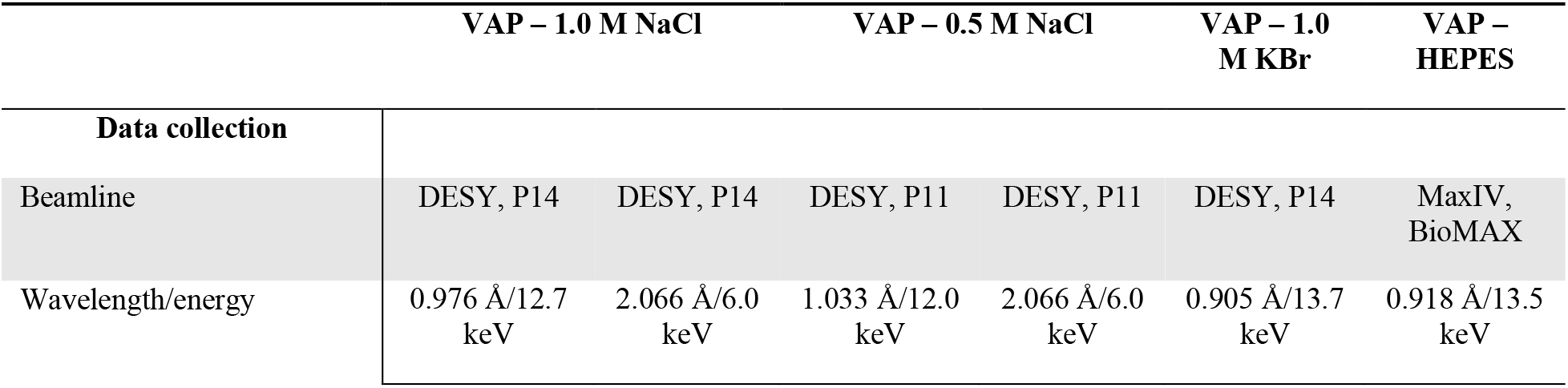
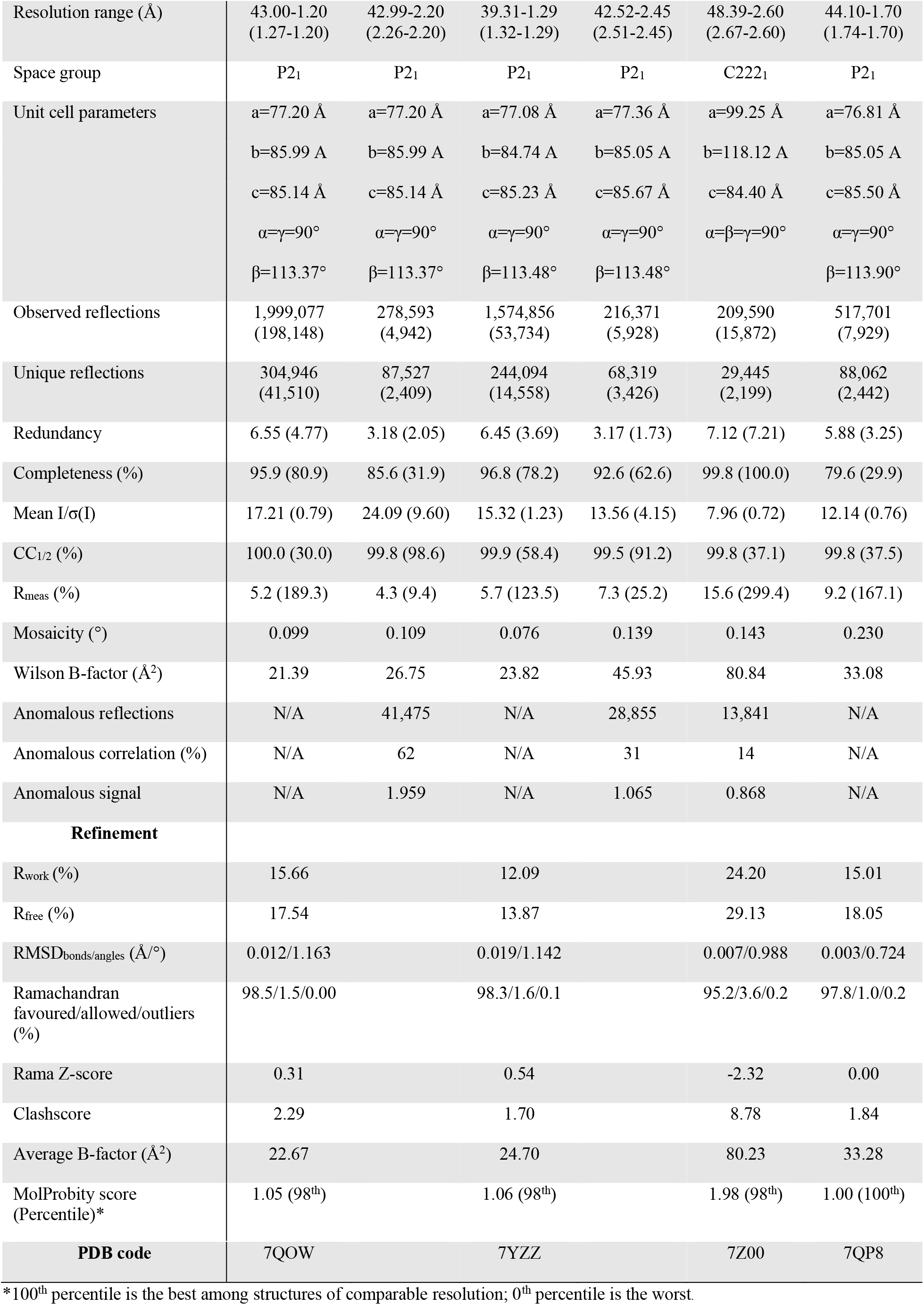
X-ray diffraction data collection and refinement statistics. Numbers in parentheses indicate statistics for the highest-resolution shell. As the low-energy (anomalous) datasets of NaCl co-crystallized VAP were only used to calculate anomalous Fourier maps, no refinement statistics are presented.

### Enzymatic activity assays

The phosphomonoester hydrolysis activity of VAP was measured using disodium *p*-nitrophenylphosphate (*p*NPP) as the substrate. Steady-state kinetics at pH 8.0 were performed in 100 mM Tris-HCl pH 8.0 and 1 mM MgCl_2_ with 0-1 mM *p*NPP, at 10 °C. The final reaction consisted of 10 μL enzyme and 990 μL reaction solution (for a final protein concentration of 8-20 μg/ml), and the formation of *p*-nitrophenol monitored for 20 s at 405 nm, using an absorption coefficient of 18.5 mM^-1^ cm^-1^. Kinetic parameters of VAP catalysis were obtained by directly fitting the data with the hyperbolic Michaelis-Menten model in GraphPad Prism (v. 8.4.2). The assays were carried out in the presence and absence of 0.5 M NaCl to estimate activation upon chloride binding. To measure activity at alkaline pH, the same assay was conducted in 100 mM CAPS pH 10.5 and 1 mM MgCl_2_.

The IC_50_ of VAP inhibition by HEPES was measured under pseudo-zero-order (the substrate concentration close to *V_max_*) hydrolyzing conditions at pH 8.0 (5 mM *p*NPP, 10 mM Tris and 1 mM MgCl_2_) and pH 10 (5 mM *p*NPP, 10 mM CAPS and 1 mM MgCl2), in the presence of 0-500 mM HEPES. To avoid alteration of pH upon addition of HEPES, its stock solution was pre-titrated to the desired pH using NaOH. A more detailed HEPES inhibition assay was carried out using the pH 8.0 steady-state assay described above, at HEPES concentrations ranging from 0-150 mM.

Enzymatic assays were made in 96-well format to increase throughput, when testing the effect of halogen salts on activity (Fig. 1A). Microplate assays were performed at 25 °C in 100 mM Tris-HCl, 1 mM MgCl_2_, 1.0 mM *p*NPP, pH 8.0 using an iD5 multimode plate reader (Molecular Devices) equipped with injectors. Enzyme assays were initiated with 5 μl injection of enzyme to a total volume of 100 μl per well resulting in 10 μg/ml final concentration of enzyme followed by shaking for 3 s, and the absorbance at 405 nm was monitored for 30 s. The enzyme stock was kept in assay buffer with the addition of 300 mM NaCl for storage stabilization between injections.

**Figure 1:**
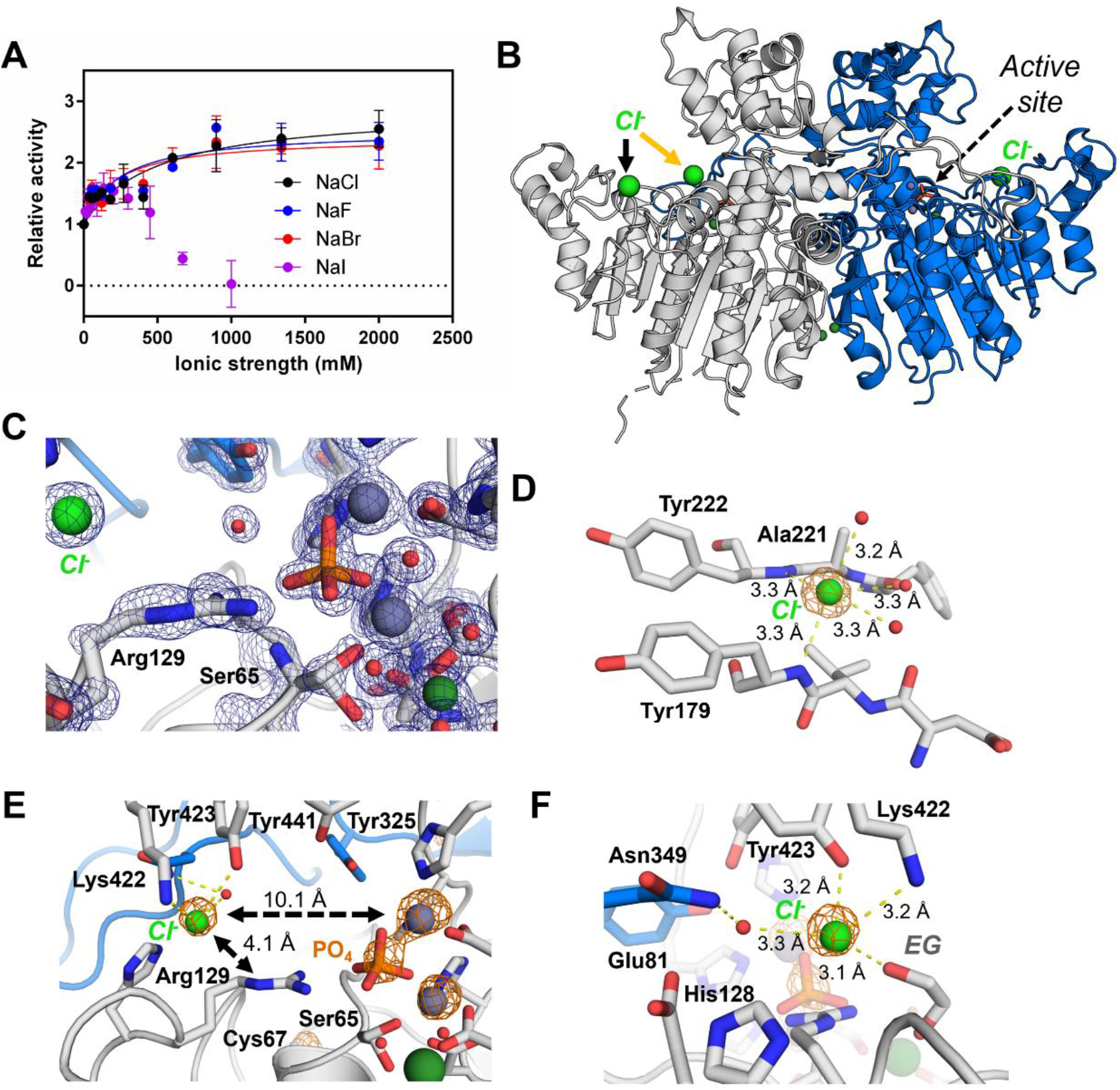
VAP binds chloride at two distinct sites. (**A**) VAP salt activation by halogen salts at pH 8.0. Activity was normalized to that at 0 M halogen salt concentration for each dataset, with error bars showing SD of the mean (n = 2) (**B**) Overall structure of the homodimeric VAP crystallised in 1.0 M NaCl. The three bound chloride ions, located by anomalous diffraction (Fig. S1), are shown as green spheres. The yellow arrow highlights the chloride ion only present in one active site, which was not observed at 0.5 M NaCl (Fig. S1). (**C**) Active site 2mF_o_-DF_c_ electron density (blue mesh at 2.0σ) of VAP in the 1.0 M NaCl crystal structure, highlighting the bound phosphate (orange/red) and chloride ion (green sphere). Active site Zn^2+^ and Mg^2+^ ions are shown as grey and dark green spheres, respectively. (**D**) Binding of chloride in the peripheral site is facilitated mainly by the backbone amides of Tyr222 and Tyr179, as well as three water molecules to complete the square bipyramidal coordination geometry. (**E**) and (**F**) show two views on the active site binding of chloride, coordinated by Lys422, Tyr423, and additionally bound by a single water molecule and ethylene glycol (EG; crystallisation cryoprotectant). Anomalous Fourier maps, derived from diffraction data collected at 2 Å (6 keV) are shown as an orange mesh contoured to 5.5σ in panels D-F. The EG molecule crystallised in proximity of Arg129 is not shown in panel E, for clarity.

### Circular dichroism spectroscopy

Circular dichroism (CD) spectroscopy was used to estimate thermal stability of wild-type VAP and mutants, as has been described elsewhere ^*39*^. In short, CD data were collected on a Jasco J-1100 instrument using 0.15-0.25 mg/mL protein sample in 10 mM borate pH 8.0, 10 mM MgCl_2_ with or without 0.5 M NaCl, in a 1-mm pathlength quartz cuvette (Hellma Analytics) at wavelengths of 260-180 nm at 25 °C.

### Thermal inactivation assay

*T_50%_*, defined here as the temperature, at which enzymatic activity was reduced to 50% of the initial activity after 30 min, was determined as previously described ^*14*^

### Small-angle X-ray scattering

SAXS experiments were carried out on the P12 beamline ^*40*^ of the PETRA III synchrotron (EMBL/DESY, Hamburg, Germany). The wavelength of the incident X-ray beam was 1.24 Å, and scattering data were collected on a Pilatus 6M detector placed 3 m from the sample, resulting in a scattering vector of 0.026-7.416 nm^-1^. Measurements were carried out at 3.2-13.8 mg/mL wild-type VAP in 20 mM Tris, 10 mM MgCl_2_, 1.0 M NaCl, pH 8.0. Measurements in lower sodium chloride concentrations proved infeasible due to sample aggregation and precipitation during measurements. Radial averaging and buffer subtraction were performed using the automated SASFLOW data analysis pipeline at the P12 beamline ^*40*^. Primary data analysis was carried out in PRIMUS ^*29*^ and distance distribution function (P(r)) analysis in GNOM ^*41*^. *Ab initio* models were generated using GASBOR ^*42*^, theoretical scattering curves of crystal structures produced in CRYSOL ^*43*^ and normal-mode analysis against SAXS data carried out in SREFLEX ^*44*^.

## Results

### Chloride binds to VAP at two distinct sites

We have previously shown that VAP is activated by monovalent anions of several types, whereby a plateau in activation was reached in the concentration range of 0.5-1.0 M ^*14*^. An identical stimulating effect on VAP activity was observed by NaCl, NaBr and NaF up to 2.0 M (Fig. 1A). NaI was inhibitory at > 0.5 M, yet showing similar activation from 0 – 0.3 M. The inhibition by NaI was most likely caused by the chaotropic nature of the iodide ion compared to the other halogens, and therefore, the observed inhibition could be a result of VAP aggregation and/or precipitation in the assay. Of these halogen salts, at these concentrations, sodium chloride is the only physiologically relevant salt in the ocean environment.

To gain detailed insight into the effect of chloride on VAP stability and activity, we set out to locate specific binding site(s) *via* crystallographic analyses. The enzyme was co-crystallized with either 0.5 M or 1.0 M NaCl, and the structures were refined at resolutions of 1.29 Å and 1.20 Å, respectively. In both cases, the structures of the homodimeric VAP were nearly identical to the previously published structure of the enzyme (PDB: 3E2D), with C_α_ root-mean-square deviations (RMSD) of less than 0.2 Å. Despite efforts towards freeing the enzymatic active sites of bound phosphate, it remained tightly bound in both active sites of the dimer. Moreover, electron density belonging to a portion of the intact C-terminal StrepTag-II affinity tag was apparent in one monomer of each dimer in the 1.0 M NaCl structure (Fig. S1). However, no inhibitory effect was noted from a synthetic peptide corresponding to the peptide tag added to kinetic assays, suggesting its presence to be a crystallization artefact (Fig. S2).

To locate bound chloride ions, the high-resolution diffraction data were supplemented with anomalous data close to the absorption K-edge of chloride, collected using an X-ray wavelength of 2.0 Å. Additional anomalous peaks corresponding to bound chloride ions were apparent in the anomalous Fourier maps (Fig. 1B and S1) in addition to peaks representing the active site Zn^2+^, phosphate ions, as well as cysteine and methionine sulfur atoms. At both 0.5 M and 1.0 M NaCl, a large anomalous peak extending to 11.1σ was apparent on the edge of the α/β-domain of the protein (in the 1.0 M co-crystal), where a solvent-exposed chloride ion was penta-coordinated in a slightly distorted square bipyramidal geometry by the backbone amides of Tyr222 and Tyr179, and three water molecules (Fig. 1D).

Along with anomalous peaks originating from the two active-site Zn^2+^ ions, and the phosphate bound in the active site in the data collected at 2.0 Å, a prominent peak, extending to 8.1σ was present in the proximity of the substrate-binding residue Arg129. This chloride ion was located 10.1 Å from the Zn^2+^ bound in metal site M1 and 4.1 Å from the N^ε^ guanidium nitrogen of Arg129. The closest residues were Lys422 and Tyr423, with distances of 3.2 Å in both cases (Fig. 1E and 1F).

It is notable that active-site chloride binding was only detected in one of the monomers in the 1.0 M NaCl structure and not seen in the 0.5 M NaCl crystal structure. This was evident in the refined crystallographic B-factors of the Lys422 in the two monomers in the 1.0 M structure, for which the average side-chain B-factor decreases from 37.8 Å^2^, in the unbound state, to 26.2 Å^2^ upon chloride binding. Increased flexibility of Lys422 in the absence of chloride binding was further noted in the lack of side-chain electron density in the subunit, which did not bind chloride.

Various other ions show anomalous characteristics at the wavelength utilized here to localize bound chloride. Calcium is a physiologically common ion that binds to various APs and has absorption characteristics like those of chloride. The assignment of non-anomalous electron density alone did not allow for distinguishing the two (Fig. 2A). Therefore, to confirm that the anomalous signal present in the NaCl co-crystal structures was in fact that of chloride, VAP was co-crystallized with another monovalent halogen salt (KBr), which has been shown to be similarly activating as chloride (Fig. 1A). Bromide has absorption characteristics different to those of chloride, with a K-absorption edge at 0.92 Å. By collecting diffraction data slightly above the bromine K-edge, where the weak anomalous signal from chloride and similarly electron-rich atoms are absent, the binding of halogen atoms could be affirmed. Crystals of VAP grown in 1.0 M KBr allowed for solving the structure at a resolution at 2.6 Å, and two distinct bromide binding sites were observed in the anomalous Fourier maps derived from the data (Fig. 2B-C, Fig. S1). Identical to that observed in the chloride co-crystals, an anomalous peak extending to 7.2σ was present on the α/β-domain, corresponding to bromide bound to VAP by the backbone amides of Tyr179 and Tyr222 (Fig. 2B). The binding mode and distances were almost identical to those observed for chloride, but due to the lower resolution of the data, water molecules coordinating the bound bromide could not be identified. Additionally, a second peak at 6.6σ was apparent in proximity of the active site chloride pocket. Although highly similar, the binding mode was not identical to that of chloride. The bromide ion bound on the positively charged patch leading up to the active site, and was primarily bound by Lys422, Arg113 and Arg153, *via* distances of 4.0-4.3 Å (Fig. 2C). Moreover, the asymmetric binding observed for the active site-bound chloride was absent in the bromide-bound structure, in which VAP crystallized in the C2221 unit cell, with a single monomer in the asymmetric unit, as opposed to the asymmetric P21 dimer observed in the chloride co-crystals.

**Figure 2:**
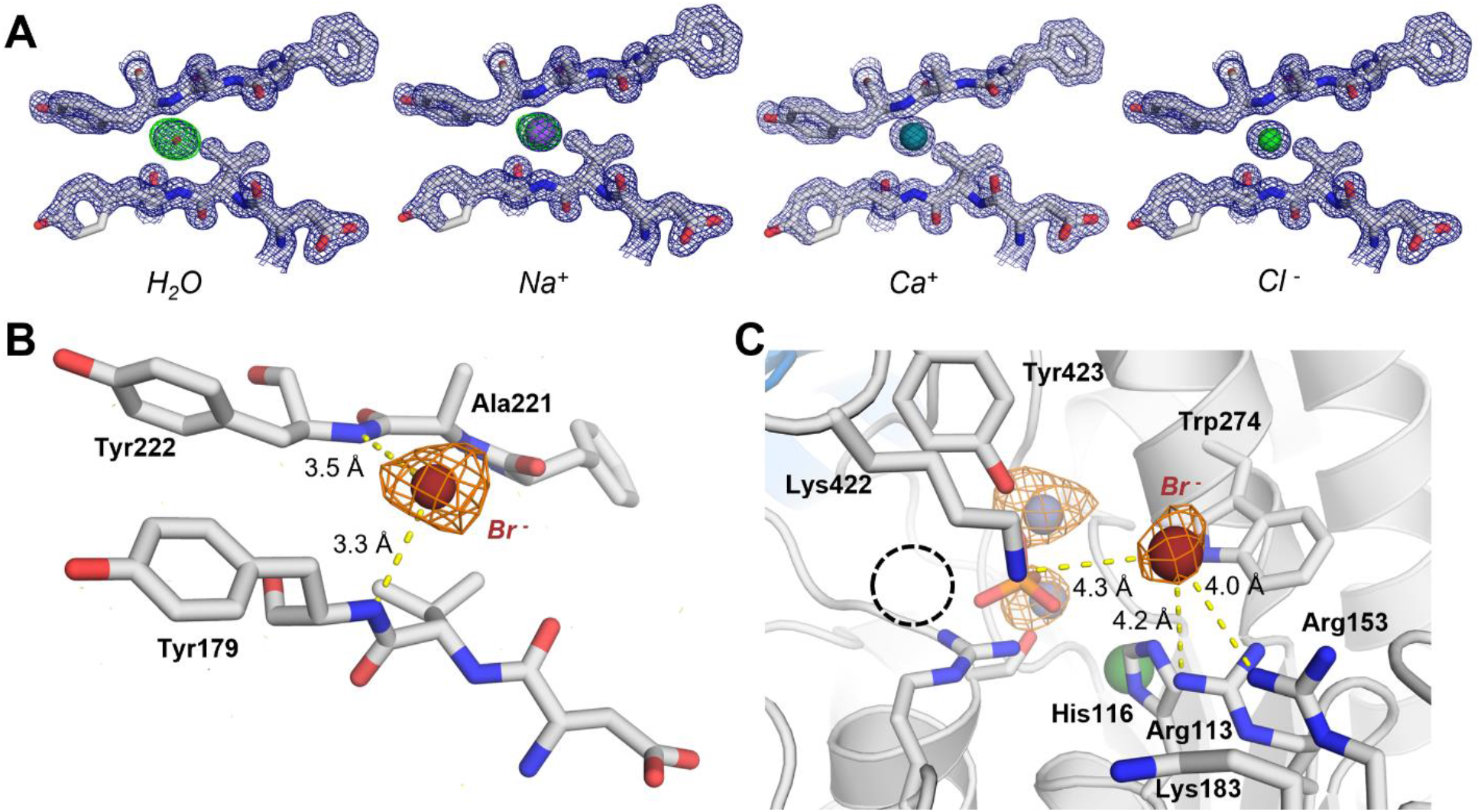
Identification of the halogen anomalous scattering ion *via* bromide co-crystallization. (**A**) Differential mapping of water and possible ions bound in the peripheral chloride binding site of the 1.0 M NaCl co-crystal structure. Placement of water or a Na^+^ ion was in strong disagreement with the data, as indicated by the large positive difference map peaks, but binding of Ca^2+^, although likely not favored by the coordination chemistry of the binding site, could not be excluded. The 2mFo-DFc electron density map is shown as a blue mesh at 2.0σ and the mFo-DFc difference map as a green/red mesh at 3.0σ. Therefore, chloride binding was confirmed through co-crystallization with bromide. (**B**) Bromide adopts a binding mode identical to chloride in the peripheral chloride binding site. (**C**) Bromide near the active site chloride pocket (highlighted with a dashed circle) was coordinated mainly by Lys422, Arg113 and Arg153. The anomalous Fourier map (from X-ray data collected at 0.9 Å) is shown as an orange mesh at 5.0σ.

Fortuitously, we found the HEPES buffer ion to be a low affinity non-competitive inhibitor of VAP with a Ki of 21.3 mM (Fig. 3A). Moreover, the inhibition appeared pH-dependent, resulting in diminished inhibition at alkaline pH (Fig. 3B). Therefore, the nature of HEPES inhibition resembled that of chloride activation, suggesting similar modes of binding. To reveal the molecular basis of the inhibition, VAP was co-crystallized with 100 mM HEPES (~5-fold *Ki*) at pH 7.0. The 1.70-Å crystal structure revealed two HEPES molecules bound in the active site of the enzyme, and as earlier, the enzyme crystallized with phosphate bound to the active site (Fig. 3C). The HEPES molecule bound at the bottom of the active site showed C-H…π interactions with Tyr441 and the Zn^2+^-coordinating His277. Additionally, the sulfonic acid moiety of the molecule was bound by Lys422 and Tyr423 at the same site as observed for chloride. Accordingly, increasing assay concentrations of chloride considerably reduced HEPES inhibition (Fig. 3D). Therefore, the inhibition mode observed for HEPES and competitive chloride binding further support active site binding of chloride, coordinated by Lys422 and Tyr423, and its relevance for activation of VAP.

**Figure 3:**
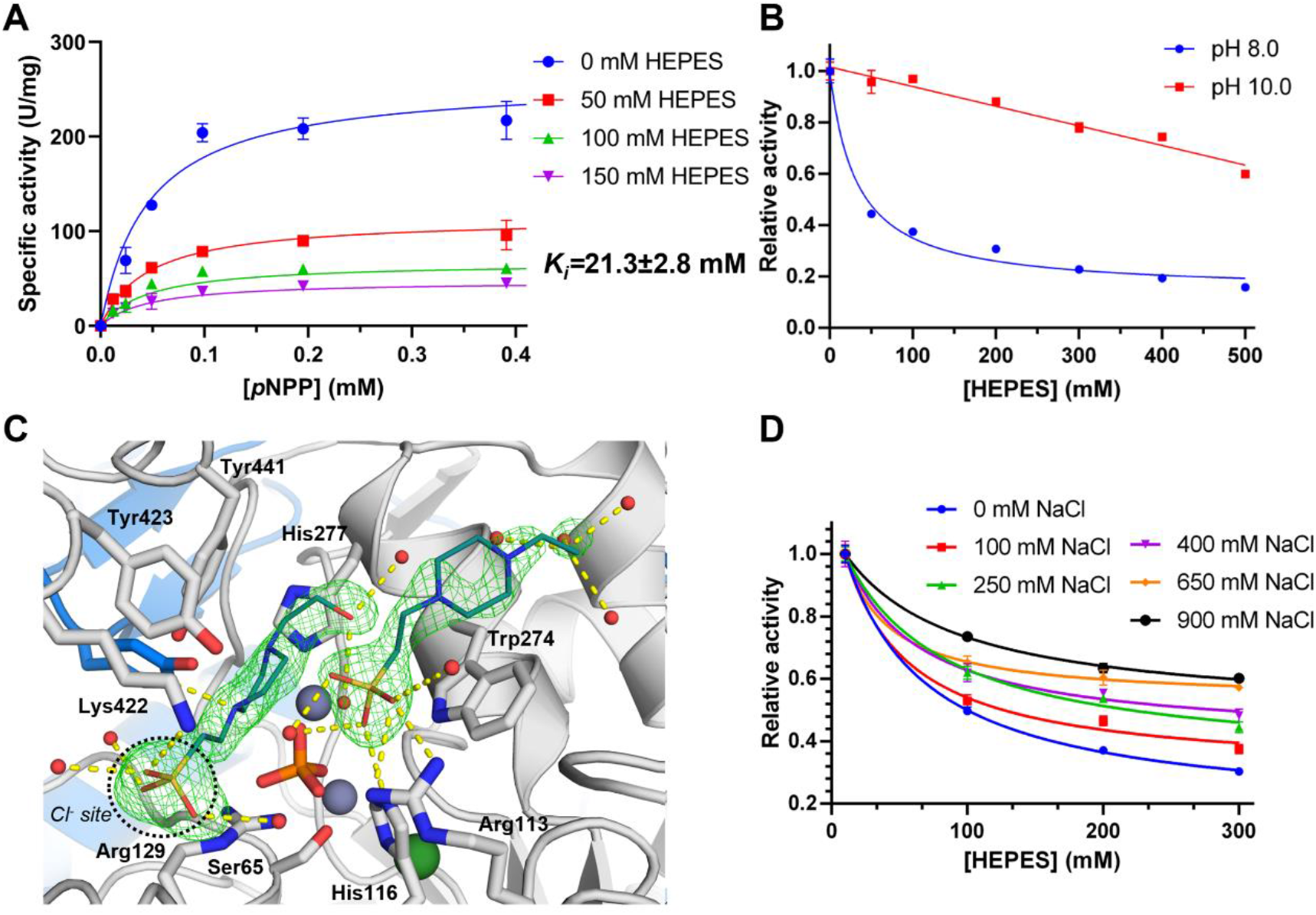
HEPES is a non-competitive inhibitor of VAP and competes with chloride for active site binding. (**A**) The *p*NPP hydrolysis activity of VAP monitored at different HEPES concentrations, at pH 8.0, shows characteristics of non-competitive inhibition. (**B**) IC_50_ plots of HEPES inhibition at pH 8.0 and pH 10 show highly pH-dependent inhibition, similarly to chloride binding. (**C**) Co-crystal structure of VAP with HEPES at pH 7.0 revealed two HEPES molecules bound in the active site of VAP. The omit mFo-Fc map is shown as a green mesh at 4.5σ. The active site chloride binding site is highlighted with a dashed circle. (**D**) Competitive binding of chloride and HEPES results in reduced inhibition at increased NaCl concentrations.

### Probing the effect of chloride binding via mutagenesis

To better understand the mechanisms underlying the stabilization and activation of VAP upon chloride binding, we set out to prevent chloride binding by mutagenesis. As binding of chloride in the peripheral site involved only backbone amides in a solvent exposed site (Fig. 1D), the aim to disrupt chloride binding by replacement of side chains was an option with an uncertain outcome. Therefore, an attempt was made to introduce a glutamate instead of the proximal Ala221 (A221D variant) to repel anionic binding in the site. Another approach was to change Tyr222 to proline (Y222P variant) to remove one of the coordinating backbone amides. These changes resulted in no significant alteration in activity or thermal stability (T_m_) of these variants (Table 2). However, a considerable decrease was measured in the active site thermal stability (*T_50%_*) of the Y222P mutant, suggesting that π-stacking between Tyr222 and Tyr179 (Fig. 1D) might be important for stabilization of the active site.

**Table 2:** Kinetic and stability parameters of VAP mutants targeting chloride binding. All measurements were carried out at pH 8.0 under hydrolyzing conditions (n=3-5 and ± SD).

| Variant | $k_{cat}$ ($s^{-1}$) | $K_M$ ($\mu M$) | $k_{cat}/K_M$ ($M^{-1} s^{-1}$) | $T_{50\%}$ ($^{\circ}C$) | $\Delta T_{50\%}$ ($^{\circ}C$) | $T_m$ ($^{\circ}C$) | $\Delta T_m$ ($^{\circ}C$) |
| --- | --- | --- | --- | --- | --- | --- | --- |
| **Control** |  |  |  |  |  |  |  |
| WT | 231±15 | 7.6±1.8 | 3.0$\cdot 10^7$ | 29.2±2.9 | -- | 51.6±0.5 | -- |
| A221D | 258±30 | 12.8±3.8 | 2.0$\cdot 10^7$ | 29.2±2.9 | -- | 51.7±0.5 | -- |
| Y222P | 194±2 | 4.9±3.4 | 4.0$\cdot 10^7$ | 23.0±0.7 | -- | 52.1±0.1 | -- |
| K422L | 165±20 | 6.6±3.1 | 2.5$\cdot 10^7$ | 30.3±3.9 | -- | 51.7±0.3 | -- |
| K422E | 49±1 | 7.0±1.6 | 7.0$\cdot 10^6$ | 24.3±0.6 | -- | 51.7±0.1 | -- |
| Y423F | 138±8 | 7.6±2.0 | 1.8$\cdot 10^7$ | 22.6±2.7 | -- | 52.2±0.2 | -- |
| K422L/Y423F | 100±12 | 8.8±2.4 | 1.1$\cdot 10^7$ | 25.9±0.8 | -- | 51.1±0.1 | -- |
| **0.5 M NaCl** |  |  |  |  |  |  |  |
| WT | 356±16 | 7.9±1.5 | 4.5$\cdot 10^7$ | 53.2±1.9 | 24.0 | 61.0±0.2 | 9.4 |
| A221D | 304±28 | 12.8±3 | 2.0$\cdot 10^7$ | 52.2±1.9 | 23.0 | 60.6±0.3 | 8.9 |
| Y222P | 328±10 | 5.1±1.9 | 6.4$\cdot 10^7$ | 52.3±1.5 | 29.3 | 60.8±0.2 | 8.7 |
| K422L | 223±19 | 7.3±2.5 | 3.1$\cdot 10^7$ | 51.7±2.1 | 21.4 | 60.7±0.2 | 9.0 |
| K422E | 88±1 | 7.3±0.1 | 1.2$\cdot 10^7$ | 52.8±1.0 | 28.5 | 60.8±0.1 | 9.1 |
| Y423F | 224±9 | 5.4±0.8 | 4.1$\cdot 10^7$ | 50.6±1.3 | 28.0 | 60.6±0.3 | 8.4 |
| K422L/Y423F | 140±2 | 11.0±2.0 | 1.3$\cdot 10^7$ | 52.1±0.5 | 26.2 | 60.2±0.1 | 9.1 |

In the active site, a chloride ion interacted with residues Lys422 and Tyr423 (Fig. 1E-F). To prevent this, the variants K422L and Y423F were made to eliminate the positive charge of Lys422 and remove the hydroxyl group of Tyr423, respectively. Additionally, the double variant K422L/Y423F was made. Lastly, an opposite charge was introduced into the chloride binding site with the K422E variant. *kcat* was reduced in all of these active site variants in the absence of chloride ions, and they showed less active site heat stability (*T_50%_*) in the absence of chloride (Table 2). These changes suggested a role for Lys422 and Tyr423 in the binding of chloride to facilitate activation of the enzyme.

No change was observed in *K_M_* values by adding 0.5 M NaCl for any of the variants, indicating that the salt did not take part in promoting or hindering the initial formation of the non-covalent enzyme-No change was substrate (E*S) complex. Therefore, chloride binding was accelerating a rate-liming step that occurs later in the reaction pathway.

### The analysis of structural dynamics of VAP suggests a basis for chloride-induced stabilization

VAP is characterized by four gene inserts. The insert that is located furthest away from the active site is insert I and contains residues 162-175 ^*16*^. Insert I forms a three-helix turn motif on the outer edge of the larger α/β-domain and is followed by a long 14-residue loop region, the attachment of which to the remainder of the enzyme is exclusively facilitated through hydrophobic interactions. This region of VAP has previously been shown by molecular dynamics (MD) simulations to be highly mobile, and its motions were suggested to modulate the activity of the enzyme ^*16*^. In the peripheral chloride binding site described here, the chloride ion bridges the loop region of insert I to the central fold of the α/β-domain, *via* binding of the amine backbones of Tyr179 and Tyr222 (Fig. 5A).

**Figure 4.**
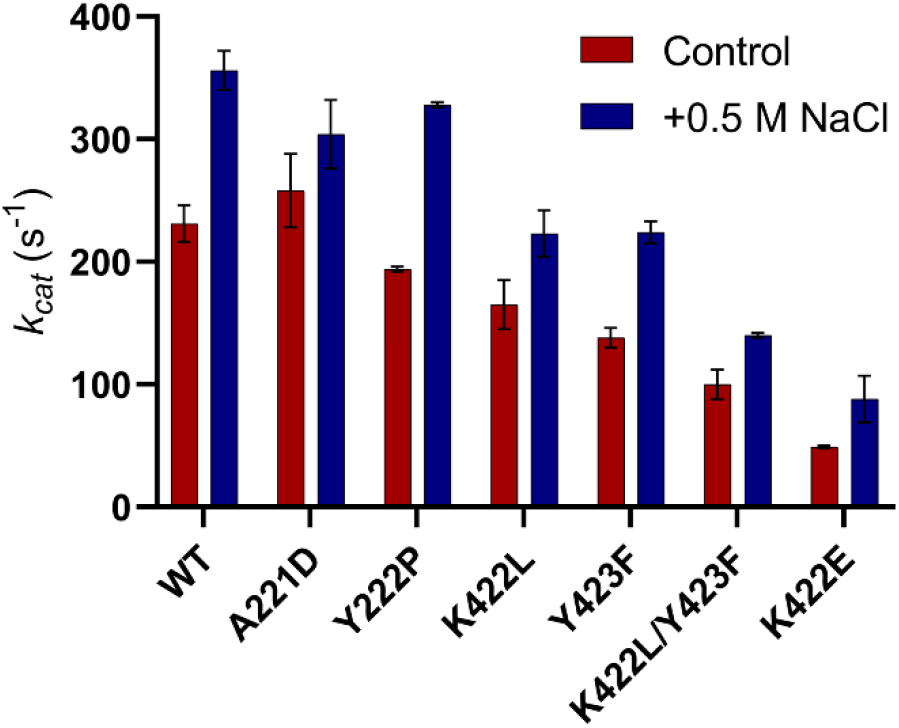
Perturbation of the chloride binding sites. Graphical depiction of data from Table 2.

**Figure 5:**
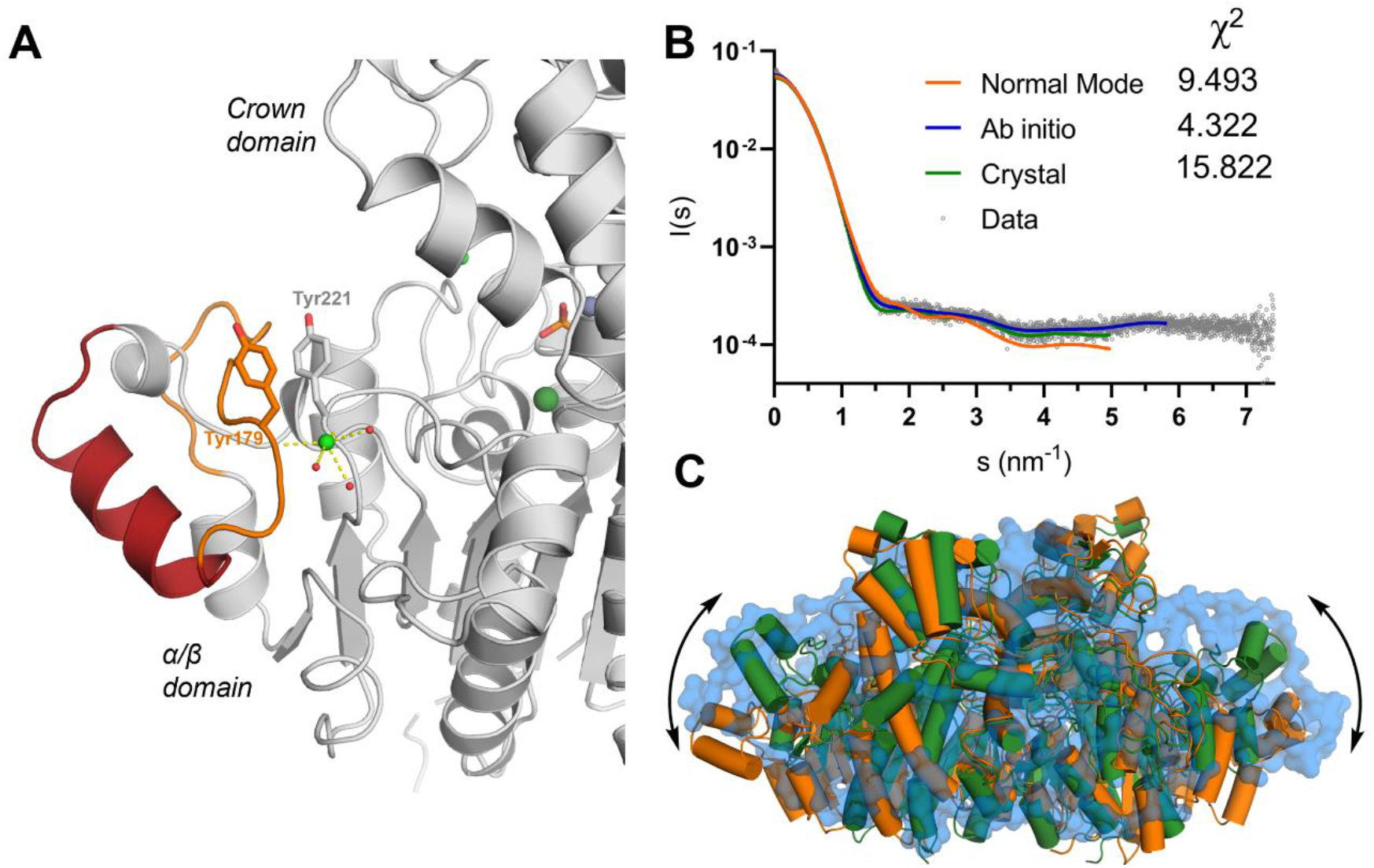
Chloride binding bridges the peripheral helix of the α/β-domain to the central fold of domain. (**A**) VAP contains a gene insert (Insert I, residues 162-175) ^*16*^ on the outer edge of the α/β-domain, colored red. Following the insert in sequence is a disordered region (residues 176-189; orange), which is bonded to the central fold of the domain via the chloride ion (green sphere) bound to Tyr179 and Tyr222. (**B**) SAXS scattering curve, measured from VAP in 1.0 M NaCl. Theoretical scattering curve was generated from the crystal structure, and fits from *ab initio* modelling in GASBOR and normal mode analysis in SREFLEX. (**C**) Superimposition of the crystal structure (green) with the best fitting model from normal mode analysis (orange) and the *ab initio* dummy chain model (transparent blue) highlights dynamic movements of VAP in solution.

We utilized SAXS to further examine the structural dynamics of VAP in solution (Fig. 5B-C, Table S1, Fig. S3). Using scattering data collected at pH 8.0 in 1.0 M NaCl, a low-resolution solution structure of VAP was obtained, and large-scale movements of the dimer in solution could be monitored. A theoretical scattering curve generated from the corresponding crystal structure of VAP showed that the solution structure differs considerably from that observed in the crystal. Normal mode analysis of the crystal structure against the SAXS data was, therefore, carried out to estimate the solution structure of the enzyme (Fig. 5B-C). The best fitting model from this analysis showed large-scale movements of the α/β-domain, most prominently in the region containing insert I, which results in a global extension of the structure and opening of the active site region. Therefore, the SAXS data gave support to the high mobility of the peripheral region of the α/β-domain previously observed in MD simulations ^*45*^. The data further suggested that rigidification of this region, via chloride binding, might strengthen the local structure to withstand initiation of unfolding, resulting in global stabilization of the enzyme.

Crystallographic B-factors can provide an estimation of the thermodynamic fluctuations of protein regions in solution with some reservations regarding packing and resolution ^*46*^. The two crystal structures of VAP obtained here, in 0.5 M and 1.0 M NaCl, had similar resolutions and almost identical unit cell parameters (Table 1), allowing comparison of crystallographic B-factors for the estimation of dynamics (Fig. 6). The B-factors of the A chain of the dimer in 0.5 M NaCl were almost identical to those observed at 1.0 M NaCl, whereas a considerable increase in the B-factors of the B chain of the 0.5 M NaCl structure was apparent compared with the 1.0 M NaCl structure. Most notably, an increase in B-factors was observed within insert I adjacent to the chloride binding site, demonstrating increased thermal fluctuations at lower chloride concentrations. Therefore, the data extracted using SAXS and comparison of crystallographic B-factors suggest that chloride binding reduced dynamic movements of this region, resulting in stabilization.

**Figure 6:**
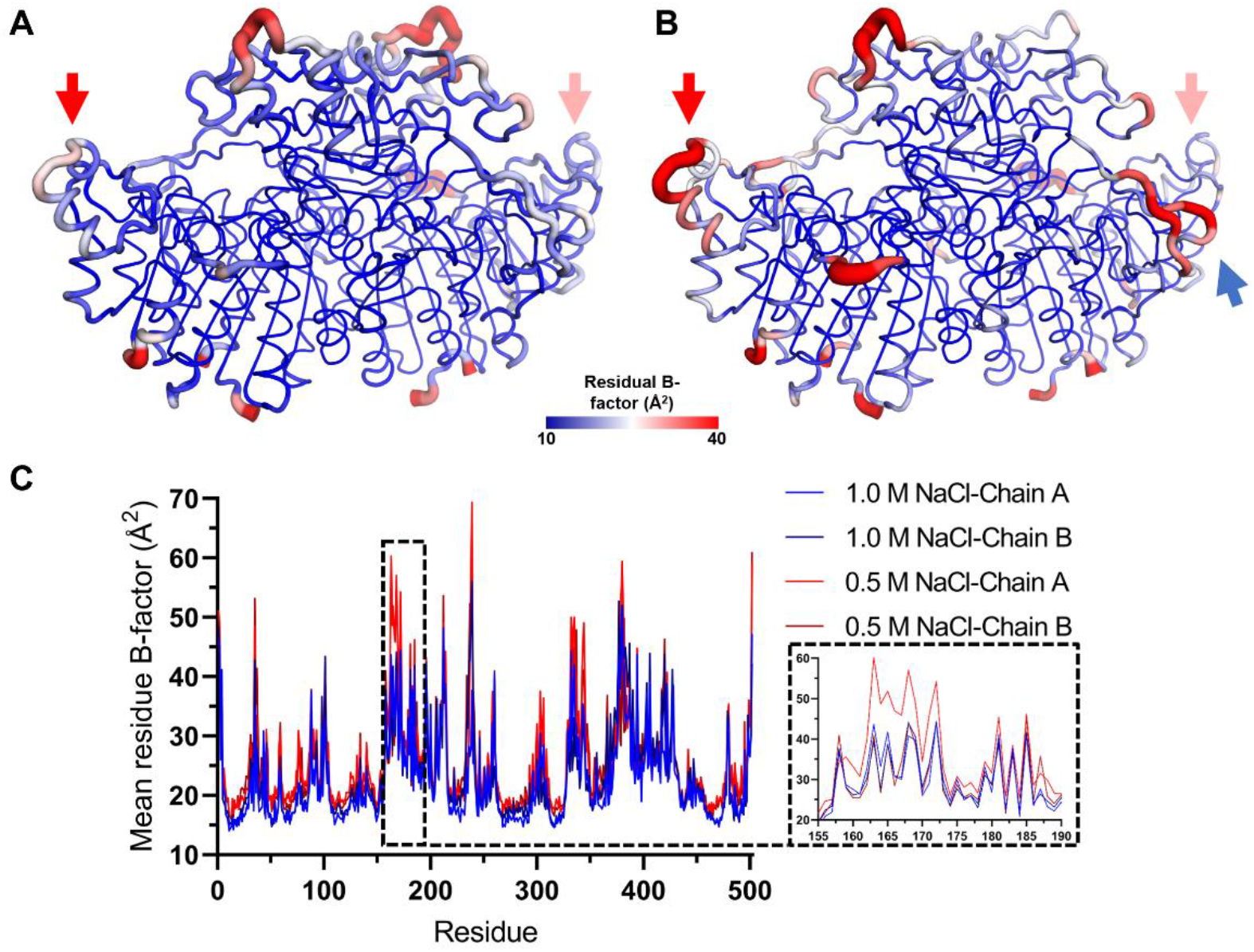
Anisotropically refined crystallographic B-factors. (**A**) VAP crystallized in 1.0 M NaCl, and (**B**) in 0.5 M NaCl suggests that increased chloride concentration reduces thermal fluctuations of the enzyme. The peripheral region of the α/β-domain, which shows higher residual B-factors at 0.5 M NaCl, is highlighted with a red arrow and the long interface loop is highlighted in the 0.5 M NaCl crystal structure in **B** with a blue arrow. (**C**) A plot showing the residual B-factors of each monomer of the two crystals structures, with an inset highlighting the more flexible region at 0.5 M NaCl highlighted in **A** and **B**.

## Discussion

The *Vibrio* alkaline phosphatase (VAP) studied here has adapted to the dual challenge of low temperature and high salinity. The presumed indicators of cold and saline adaptation are generalized as increased structural flexibility and a low isoelectric point, respectively ^*47, 48*^. As salt concentration increases, the influence on the physical properties of water and ionic interactions becomes more telling. Some ion types support the regular structural interactions in water that drive the hydrophobic effect, while others allow more facile mixing of water molecules with non-polar solutes by reducing that adhesion between water molecules. Ion pairing with the enzyme and reagents will also become more intensive as the concentration rises. Thus, solvation and dissolvation of reagents may give rise to non-specific rate enhancements. In this respect, sodium and chloride are relatively inert salts compared with other ions in the Hofmeister series. Therefore, the effects observed here may be more specifically linked with binding to sites in the enzyme structure. Notably, NaCl was ineffective as an activator of VAP at high pH (>10), and the same was observed before with *V. alginolyticus* AP ^*17*^. As the pH-dependent effect was traced to the halogen ions in particular, it is likely that binding is to positively charged site that carries a dissociating proton(s). The groups found in proteins with pKa values above 9 are mainly Lys, Arg and Tyr, but may possibly include other groups where the pKa is non-standard, such as His ^*49*^. The water molecules bound to the metal ions could also be affected by pH. Here, the specificity of the activation is only limited to the anionic character of the ions, as other anions had a stimulating effect on VAP similar to chloride (Figure 1 and ^*14*^). High ionic strength was previously postulated to affect the equilibrium between two enzyme conformations of *E. coli* AP differing in catalytic efficiency ^*25, 26*^, but it remains uncertain what steps in the reaction pathway are actually modified by NaCl.

Earlier, we suggested - based on precedence from others - that the chloride-induced rate increase in VAP was due to competition with the phosphate product for attraction to the active site metal ion core. This would facilitate an acceleration of the rate-limiting step of catalysis at neutral pH, the product release ^*14*^. However, the presence of phosphate in both active sites of the crystal structures observed in the present work, even at 1.0 M NaCl, seems to highlight the inability of chloride to displace phosphate. Another effect of chloride might be to reduce the positive electrostatic potential in the active site, due to the three metal ions and positively charged side chains. Thus, chloride binding could transiently affect the electrostatic potential and mobility within the active site. This could facilitate product dissociation and substrate binding. A mechanism for product release has been suggested for *E. coli* AP (ECAP), based on the observed torsion-angle mobility of Arg166 (Arg129 in VAP). This residue binds the monophosphoester substrate and provides stabilization of the charged intermediates along the reaction pathway through interaction of the guanidium group with two of the phosphate oxygens. Arg166 has been observed in an outwards-facing conformation, in the near inactive S102T mutant, which has been suggested to facilitate product release and substrate binding ^*50*^. The additional methyl group introduced when Ser was mutated to Thr in ECAP resulted in steric hindrance, which inhibited the interaction of Arg166 to the bound phosphate, suggesting that the Arg residue might adopt this outward-facing conformation in the unbound state (see Fig. 7A-B). Furthermore, similar conformational freedom of the substrate-binding Arg has been observed in crystal structures of shrimp AP ^*51*^, and rat intestinal AP ^*52*^, further suggesting its functional relevance, and this can be modelled into VAP as shown here (Fig. 7C-D). The position of chloride in the active site would place it within 4 Å of the guanidium group of the flipped Arg129 (Fig. 1E-F), possibly facilitating its stabilization and thereby the rate of product release. Based on this observation, a revised reaction scheme, which accounts for chloride activation, can be presented for VAP (Fig. 7E). In this scheme, the outwards facing (product-releasing) rotamer of Arg129 (in E*) is likely stabilized by binding of chloride to Lys422 and Tyr423, resulting in accelerated product release and overall rate enhancement. The crystal structure of VAP in 1.0 M NaCl showed the E·P complex, with likely a small population of Cl^-^·E*+P, resulting in the observed partial occupancy of Cl^-^ in the active site chloride pocket. However, whether this conformation of Arg129 does appear in the apo chloride bound state of the enzyme could not be determined, as our efforts to dissociate the bound phosphate were unsuccessful.

**Figure 7:**
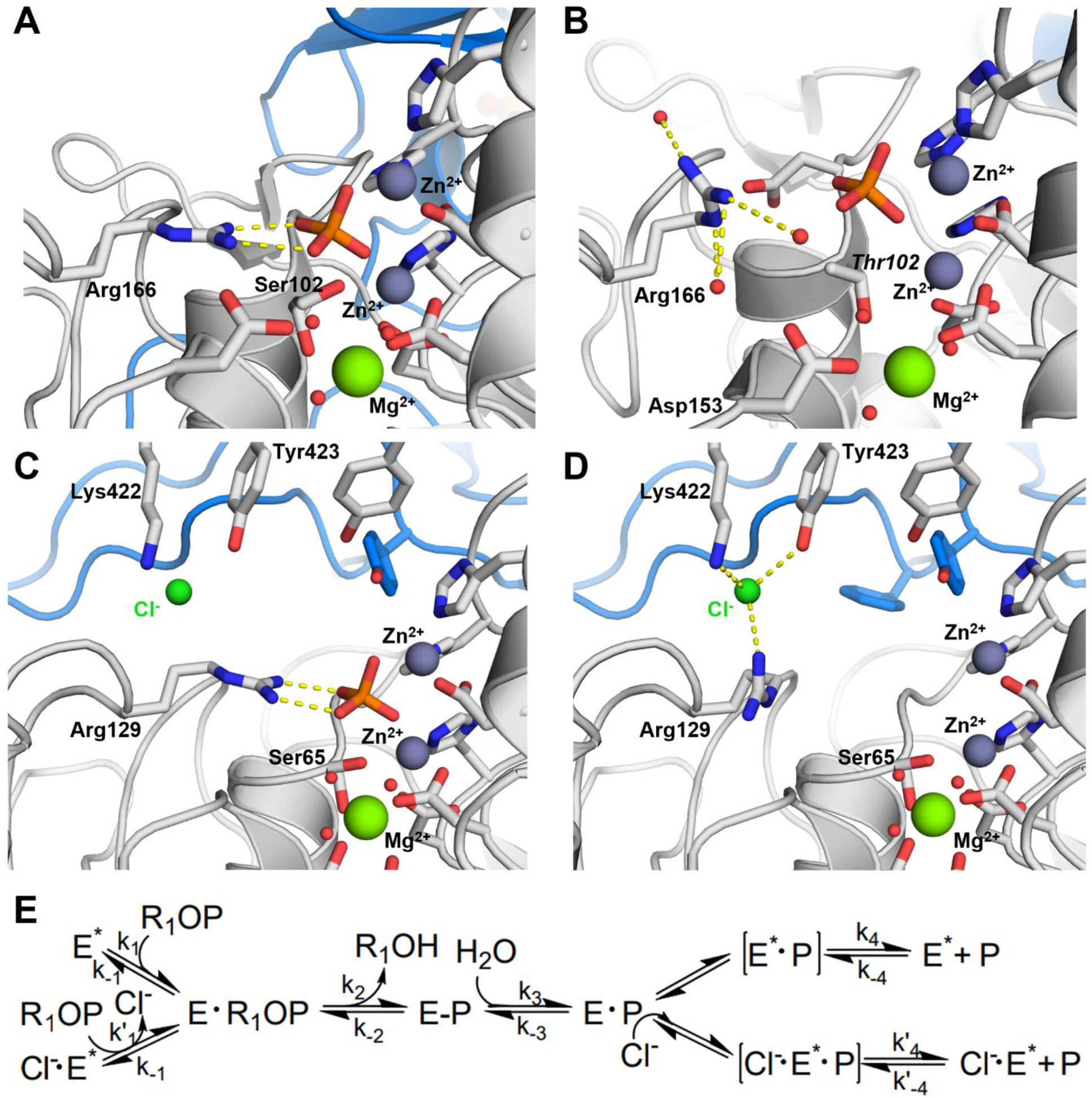
A scheme for chloride activation *via* rotational stabilization of Arg129. (**A**) E. coli AP crystallized in the E·P state, where Arg166 directly binds the enzymatic reaction product (PDB: 1ED8) ^*53*^ and (**B**) with the substrate-binding Arg166 in the outwards-facing conformation, in the phosphate-bound S102T mutant (PDB: 2G9Y,) ^*50*^. VAP was crystallized here in the E·P state with bound chloride as shown in (**C**). The positioning of chloride in the crystal structure suggests that it stabilizes the hypothetical outwards-facing conformation of Arg129 in VAP, in the absence of phosphate, to facilitate rate enhancement, illustrated in (**D**). (**E**)The proposed reaction scheme of VAP catalysis, accounting for chloride rate enhancement of the rate limiting step, where *k’_4_*>*k4*.

Few examples exist in the literature describing the effect of chloride ions on enzyme function and structural detail. Chloride activation of α-amylase and angiotensin converting enzyme (ACE) was suggested to involve: (i) positioning of catalytic residues to stabilize the enzyme-substrate complex and (ii) fine tuning of pK_a_ values of catalytically important residues ^*54*^. Removal of Cl^-^ by dialysis or gel filtration fully inactivated chloride-dependent α-amylases, but activity was recovered by addition of Cl^-^ or Br^-^ ^*55*^. We have also observed that after unfolding of VAP in urea, reactivation could only be observed upon addition of fresh buffer containing 0.5 M NaCl ^*39*^. Chloride-dependent pancreatic α-amylase has a conserved Cl^-^ pocket consisting of two Arg residues, an Asn residue and a water molecule ^*56*^. In *A. haloplanctis* amylase, one of the Arg residues is replaced by Lys ^*55*^. Pokhrel et al. noticed similarities between the activation of α-amylase and ACE, whereby removal of chloride triggered formation of a salt bridge of either Arg or Lys in proximity of the chloride binding pockets, and proposed a similar mechanism for chloride binding in photosystem II (PSII) ^*54*^. For VAP, the active site chloride binding pocket has no acidic residues in the proximity of Lys422. Thus, it is not possible to speculate on salt bridge disruption as part of the catalytic activation by chloride. However, the substrate-binding guanidium group of Arg129 is only 4.1 Å away from the chloride ion, which might provide the opportunity for chloride binding to alter the pKa of the side chain for the modulation of substrate and product binding. ACE has shown complex chloride regulation of activity, and two binding sites for chloride exist ^*57*^. Mutational studies showed that the chloride 1 pocket has limited impact on activity but was believed to stabilize the binding of substrate subsites. The chloride 2 pocket was shown to mediate both substrate binding and activity. The chloride 2 pocket had very similar binding modes, as with the active site chloride in VAP, coordinated by Arg and Tyr (Lys and Tyr in VAP). Interestingly, the distance between the catalytically active Zn^2+^ and chloride ions in ACE and VAP is very similar (~10 Å). The maximal activity for ACE plateaued at a lower concentration of chloride than VAP, yet it was substrate- and pH-dependent ^*58*^. Moreover, ACE was not selectively activated by chloride, but other halogens as well as nitrate were also effective, depending on the substrate used. The substrate dependence of chloride activation of VAP needs to be further tested to see if these similarities hold.

Recent single-molecule experiments ^*49, 59–63*^ have generally shown two active states of APs from several different origins. Earlier studies using capillary electrophoresis to see product formation by individual AP molecules indicated up to ten differently active kinetic states, with three of the active forms being dominating ^*59*^. All these studies have used synthetic substrates to monitor the rate of formation of the initial product from the first reaction step in a multi-step reaction mechanism, which is not considered rate-determining at alkaline pH. Thus, subsequent steps need to be considered for the full picture. Furthermore, the effect of dimer dissociation in these experiments can be difficult to take into account. We have previously reported VAP dimers to readily dissociate upon dilution to < 1 nM ^*14*^, but single molecular experiments are generally conducted close to fM concentrations. Another approach was recently described that does not depend on measuring reaction product(s) to monitor protein conformational changes but involved the use of a nanoantenna strategy to monitor structural changes of calf intestinal AP during catalysis. It was concluded that the dye-enzyme interaction mediated by the nanoantenna allowed monitoring of the conformational changes on the enzyme surface during its function.

Five distinct conformational states were characterized, including the transient enzyme–substrate complex ^*64*^.

Unfortunately, methods for studying the fast dynamics of small structural changes associated with kinetic pathways in enzymes are limited, and they often involve attaching probes to the enzyme. For APs, it is becoming clearer that several structural states exist with different kinetic efficiencies. These states do interchange in harmony with changes in several solvent properties, such as pH, salt, inhibitors, or kaotropic salts, and physical conditions, such as temperature. Further development of sensitive single-molecule techniques combined with classical methods will undoubtedly link known functional effects to specific structural alterations in APs, distinguishing between the various enzyme, enzyme–substrate and enzyme-product forms.

## Conclusions

The present work identifies two chloride binding sites in VAP. One is located deep in the active site, close to where the substrate phosphoryl group binds, while the other is on the periphery, away from the dimer interface and active sites. The chloride ion activation observed in wild-type VAP was still present after key residues in either site had been substituted with the aim to prohibit chloride binding. Despite one of the chloride binding sites being close to the phosphoryl-group binding site, it did not seem to promote enhanced product release by exchanging with the inorganic phosphate product. The way in which the entry of chloride ion(s) into the large open active site of VAP affects catalysis appears to be a more complex interplay of effects than anticipated. The abundance of chloride ions could both modulate local fine tuning of the electrostatic potential gradients in the active site ^*65, 66*^ as well as directly affect protein dynamics in the area and promote necessary interactions and cooperation between the subunits.

Chloride binding to the second peripheral site distal to the active site resulted in a large reduction in mobility and enhanced heat stability. Yet, the decrease in enzyme dynamics in this local area at high NaCl concentrations did not prevent a large chloride-induced increase in enzyme activity observed under the same conditions. It is suggested that chloride ions may facilitate a rate-limiting conformational change, possibly by regulating the movement of Arg129, a key substrate/product-binding residue. Taken together, our results highlight the complex role that ions in the protein medium can have, and how NaCl can simultaneously provide stabilization and increase catalytic efficiency of a cold-active enzyme adapted to marine salt conditions.

## ASSOCIATED CONTENT

### Accession codes

VAP (UNIP: Q93P54)

Crystal structure coordinates and experimental data are available under protein data bank (PDB) accession codes 7QOW, 7YZZ, 7Z00 and 7QP8.

X-ray crystallography raw data and anomalous maps are located upon request at:

1.0 M NaCl remote data: doi.org/10.5281/zenodo.5807483

1.0 M NaCl anomalous data: doi.org/10.5281/zenodo.5809400

0.5 M NaCl remote data: doi.org/10.5281/zenodo.5878920

0.5 M anomalous data: doi.org/10.5281/zenodo.6857326

KBr anomalous + data: doi.org/10.5281/zenodo.6857655

HEPES remote data: doi.org/10.5281/zenodo.5812840

## AUTHOR INFORMATION

### Author Contributions

SM and JGH planned experiments; SM, JGH and BÁ performed experiments; SM, JGH, BÁ and PK analyzed data; BÁ and PK contributed reagents and access to facilities; SM, JGH and BÁ wrote the paper; All authors took part in revision of the manuscript draft.

## Acknowledgments

We want to thank Dr. Bjarte Aarmo Lund at the department of Chemistry, arctic University of Tromsø for assistance with initial screening and data collection at the BioMax beamline. We acknowledge MAX IV Laboratory for time on the BioMAX Beamline under Proposal 20190447. Research conducted at MAX IV, a Swedish national user facility, is supported by the Swedish Research council under contract 2018-07152, the Swedish Governmental Agency for Innovation Systems under contract 2018-04969, and Formas under contract 2019-02496. Parts of this research were carried out on beamline P11 at DESY, a member of the Helmholtz Association (HGF). Furthermore, we wish to thank EMBL/DESY for access to beamlines P12 and P14.

We acknowledge the use of the Core Facility for Biophysics, Structural Biology, and Screening (BiSS) at the University of Bergen, which has received infrastructure funding from the Research Council of Norway (RCN) through NORCRYST (grant number 245828) and NOR-OPENSCREEN (grant number 245922).

## Abbreviations

AP: alkaline phosphatase
VAP: Vibrio sp. alkaline phosphatase
SAXS: small-angle X-ray scattering
ACE: angiotensin converting enzymes
ECAP: *E. coli* alkaline phosphatase
IPTG: isopropyl β-D-1-thiogalactopyranoside
pNPP: para-nitrophenyl phosphate
MWCO: molecular weight cut-off
MR: molecular replacement
CD: circular dichroism
RMSD: root mean square deviation
EG: ethylene glycol
PSII: photosystem II

