## Supplementary data for "Structural characterization of functionally important chloride binding sites in the marine *Vibrio* alkaline phosphatase"

### Figure S1

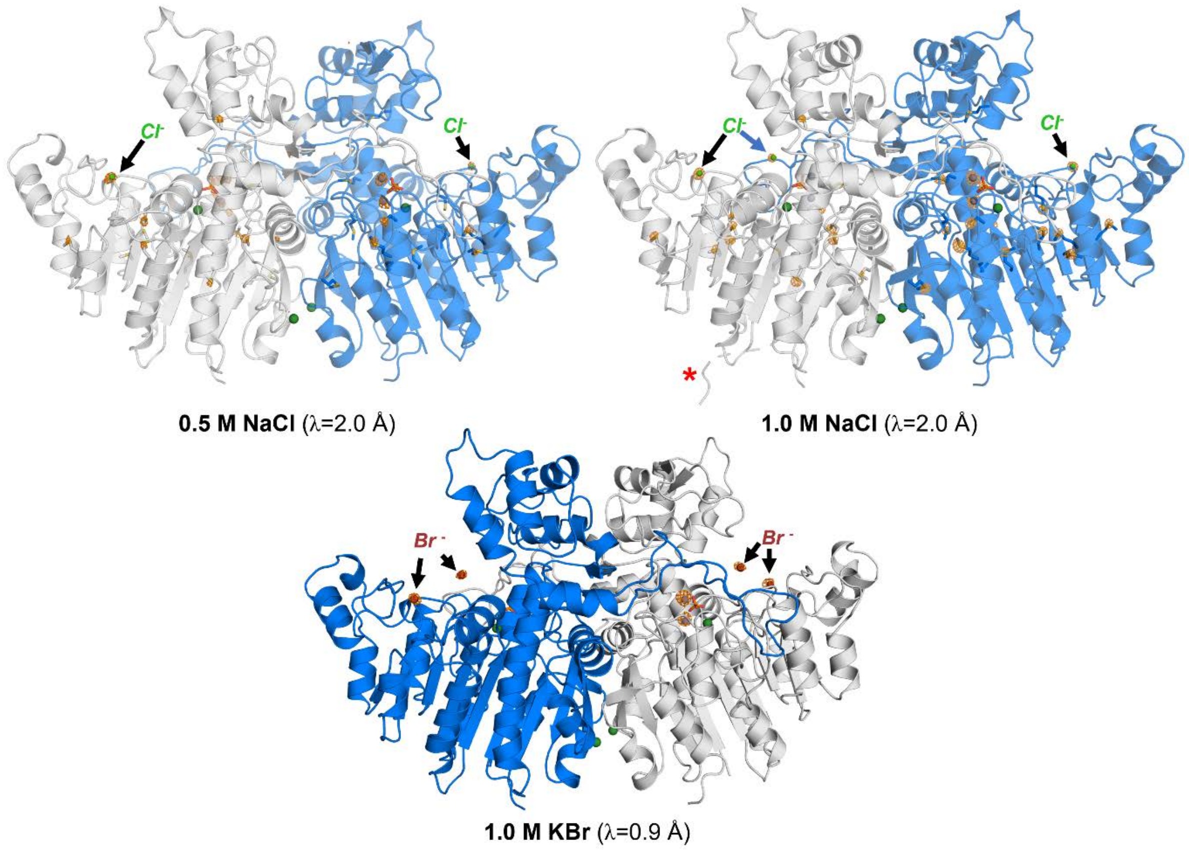


**Figure S1:** Overview of the anomalous maps used to locate chloride and bromide ions bound to VAP. In all cases, the derived anomalous Fourier maps are shown as an orange mesh contoured to 5.5σ (chloride co-crystals) and 5.0σ (bromide co-crystal). The X-ray wavelength used for collection of the anomalous data is indicated in parentheses. In all cases, additional anomalous signal arose from the active site $Zn^{2+}$ ions and at 2.0 Å additional signals appear from sulfur (in cysteine and methionine residues) and the phosphorus core of the bound phosphate.

### Figure S2

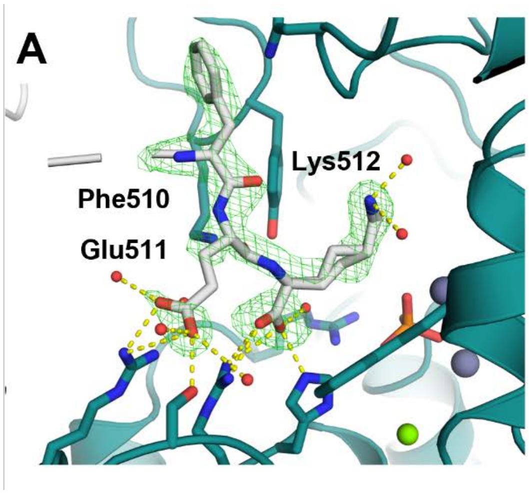


B

| Sample | U/mg |
|---|---|
| Control, 5 mM *p*NPP | 515.6±1.8 |
| +StrepTag (50 μM) | 520.7±6.0 |
| Control, 0.5 mM *p*NPP | 207.0+7.5 |
| +StrepTag (50 μM) | 205.1±2.5 |
| Control, 0.05 mM *p*NPP | 23.3±0.7 |
| +StrepTag (50 μM) | 20.5±1.6 |

**Figure S2:** Active site binding of phosphate and the C-terminal StrepTag-II is a crystallization artifact. (**A)** Active site binding of the C-terminal StrepTag-II affinity purification tag in the 1.0 M NaCl co-crystal. The tag from chain B of the asymmetry dimer stretched into the active site of the adjacent symmetry related mate to bind in its active site. A Fo-Fc omit map is shown at 2.0σ as a green mesh. (**B)** Obtained steady-state rates of *p*NPP hydrolysis by VAP in the absence and presence of 50 µM of a StrepTag peptide (Trp-Ser-His-Pro-Gln-Phe-Glu-Lys), demonstrating no apparent inhibition. Rates shown in B were measured in 100 mM Tris, 1 mM $MgCl_2$, pH 8.0 at 25°C. Final concentration of VAP in the assay was 5.85 ng/ml (~1 nM) (n=4).

### Figure S3

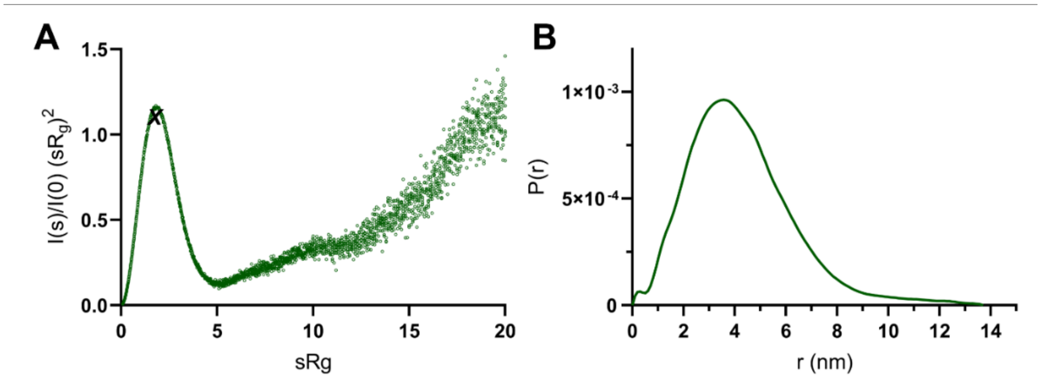

**Figure S3:** Additional processing of the SAXS curve measured from VAP in 1.0 M NaCl (Fig. 6B). **A** Dimensionless Kratky plot. The maximum of an ideal, spherical particle ($\sqrt{3}, 1.104$) is marked by ***X***. **B** Distance distribution profile.

**Table S1:** Size and shape parameters of VAP derived from SAXS data collected at 1.0 M NaCl (Shown in Fig. 6B).

| $I_0$ | $R_g$ (nm) | $D_{max}$ (nm) | Porod vol. ($nm^3$) | $Q_P$ Mw (kDa) | Monomer Mw (kDa) | Oligomeric state |
|---|---|---|---|---|---|---|
| 0.055 | 3.24 | 13.65 | 161.2 | 96.7 | 58.5 | Dimer |
